# Crosstalk between β-catenin and WT1 signalling activity in acute myeloid leukemia

**DOI:** 10.1101/2021.11.06.467095

**Authors:** M Wagstaff, O Tsaponina, G Caalim, H Greenfield, L Milton-Harris, EJ Mancini, A Blair, KJ Heesom, A Tonks, RL Darley, SG Roberts, RG Morgan

## Abstract

Wnt signalling is an evolutionary conserved signal transduction pathway heavily implicated in normal development and disease. The central mediator of this pathway, β-catenin, is frequently overexpressed, mislocalised and overactive in acute myeloid leukaemia (AML) where it mediates the establishment, maintenance and drug resistance of leukaemia stem cells. Critical to the stability, localisation and activity of β-catenin are the protein-protein interactions it forms, yet these are poorly defined in AML. We recently performed the first β-catenin interactome study in blood cells of any kind and identified a plethora of novel interacting partners. This study shows for the first time that β-catenin interacts with Wilms tumour protein (WT1), a protein frequently overexpressed and mutated in AML, in both myeloid cell lines and also primary AML samples. We demonstrate crosstalk between the signalling activity of these two proteins in myeloid cells, and show that modulation of either protein can affect expression of the other. Finally, we demonstrate that WT1 mutations frequently observed in AML can increase stabilise β-catenin and augment Wnt signalling output. This study has uncovered new context-dependent molecular interactions for β-catenin which could inform future therapeutic strategies to target this dysregulated molecule in AML.

**Article summary:**

- β-Catenin is frequently dysregulated in acute myeloid leukemia (AML) and protein interactions govern its stability, localization and activity, but these are poorly defined in AML.
- This study shows for the first time that β-catenin and Wilms tumour protein (WT1) interact and influence each other’s expression level and signalling activity in AML cells, which could inform future therapeutic strategies.

## Dear Editor

Acute myeloid leukemia (AML) affects around 3,200 people annually in the UK (Cancer Research UK statistics, accessed May 2021) and is a significant health burden. New targeted therapies in AML are showing promising efficacy but further novel treatments are required that target specific molecular aberrations, reduce toxicity, and induce long lasting remissions. One such molecular target of considerable interest given it’s frequent dysregulation in AML is Wnt/β-catenin signaling.^1^

β-Catenin is the central mediator of Wnt signaling and frequently overexpressed in AML where it is associated with poor prognosis.^2^ Wnt/β-catenin is also known to drive the emergence and maintence of leukemia stem cells in AML.^3^ Protein interactions are critical to the stability, localisation and activity of β-catenin, and we recently performed the first protemeomic analyses of the β-catenin interactome in myeloid cells.^4^ This study identified Wilms tumour protein (WT1) as a putative novel interaction partner in myeloid cells.^4^ WT1 is also overexpressed and mutated in AML where it confers inferior survival,^5,6^ yet the interplay between these two signalling proteins has not been examined previously within a hematopoietic context.

In order to idenitfy appropriate cell lines in which to study β-catenin:WT1 interplay we first performed a screen of myeloid cell lines to examine β-catenin and WT1 protein expression. We observed a statistically significant correlation between β-catenin and WT1 expression across 16 myeloid cell lines, with 50% (8/16) co-expressing β-catenin and WT1 to varying degrees (Figure 1A and B). There was no particular association of this correlation with cell line morphology or genotype. To validate the interaction between β-catenin and WT1 we perfomed the reciprocal WT1 co-immunoprecipitation (Co-IP) in one of the β-catenin/WT1 co-expressing cell lines (HEL) and confirmed interaction under basal (DMSO or PBS/BSA), pharmacologically activated (CHIR99021; GSK3β inhibitor), and naturally activated (recombinant rWNT3A) Wnt signalling conditions (Figure 1C). We further validated interaction in two other myeloid cell lines (KG-1 and K562; Supplementary Figure S1A) and confirmed this interaction also occurs in the nucleus (Figure 1D). WT1 is an RNA-binding protein (RBP),^7^ and β-catenin has also been shown to bind RNA,^8^ so to confirm this interaction was not indirect via RNA binding we repeated WT1 Co-IPs (+/-CHIR99021) after first confirming complete digestion of RNA through RNase pre-treatment of cell lysates (Figure 1E). As shown in Figures 1F, G and H, the β-catenin:WT1 interaction remained in K562, HEL and KG-1 cells under both basal (DMSO) and stimulated (CHIR99021) Wnt signalling. We further wanted to ascertain whether this protein interaction was direct using recombinant versions of purified β-catenin and WT1 protein but failed to detect association (Supplemental Figure S1B and C). This suggested that perhaps the interaction is mediated through a post-translational modification, common partners such as WTX,^9,10^ or cellular structures like DNA, given their well documented roles in transcription.

**Figure 1.**
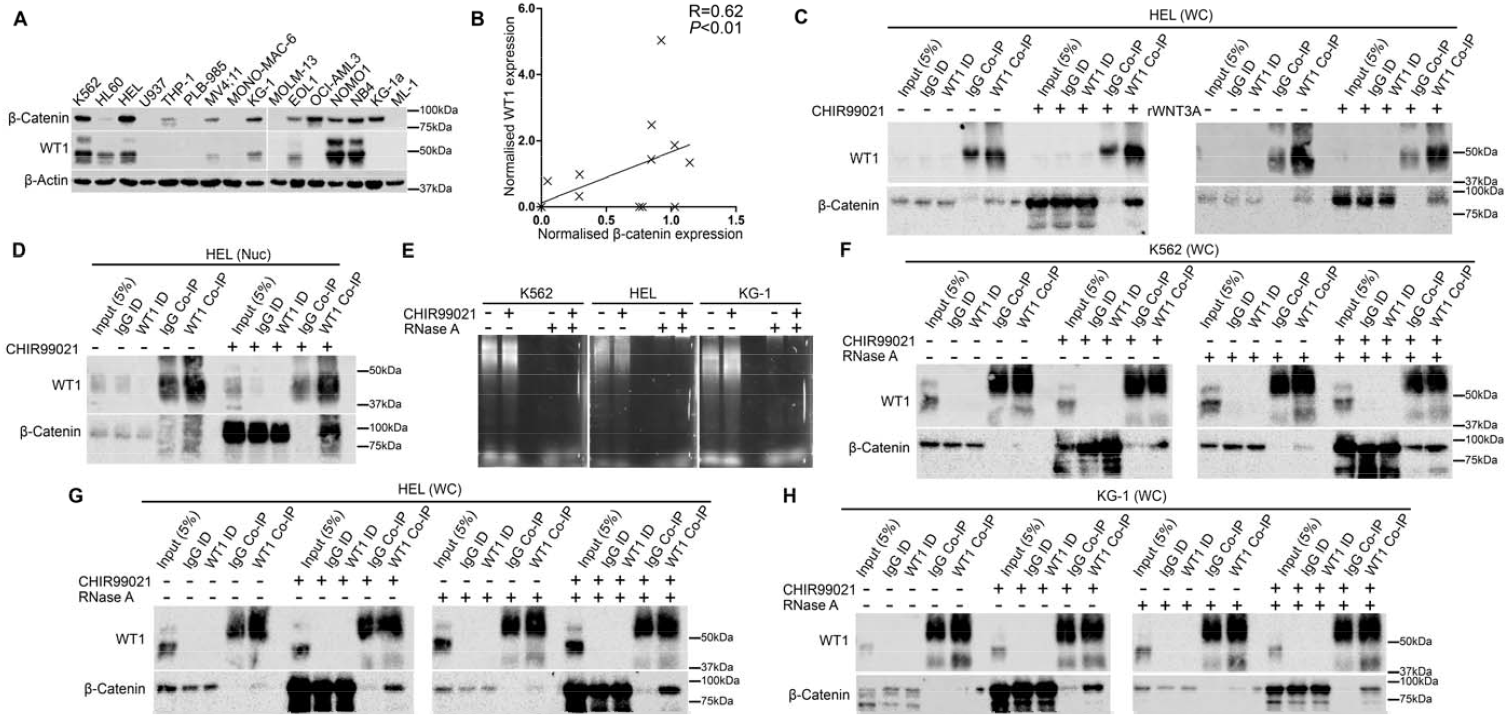
Association between β-Catenin and WT1 in myeloid cell lines and primary AML samples. (**A**) Immunoblot of myeloid leukemia cell lines showing the relative level of β-catenin (∼92kDa) and WT1 (∼50kDa) protein, with β-actin used to assess protein loading. (**B**) Summary scatter plot showing the correlation (Spearman Rank R=0.62, P<0.01) between relative β-catenin and WT1 protein expression in myeloid cell lines (normalised to β-actin expression within the cell line). (**C**) Immunoblots showing the level of β-catenin protein present in WT1 Co-IPs derived from HEL whole cell (WC) lysate under basal (DMSO), pharmacologically induced (5µM CHIR99021) and naturally induced (1µg/mL rWNT3A) Wnt signalling conditions. ID= immunodepleted lysate. (**D**) Immunoblot showing the level of β-catenin protein present in WT1 Co-IPs derived from HEL nuclear (Nuc) lysate under basal (DMSO) versus induced (5µM CHIR99021) Wnt signalling conditions. (**E**) Agarose gel electrophoresis showing the stability of total RNA in K562, HEL and KG-1 cell lysates +/-5µM CHIR99021 treated overnight +/-20µg/mL RNaseA prior to WT1 Co-IP analysis. Immunoblots showing the level of β-catenin protein present in WT1 Co-IPs derived from (**F**) K562, (**G**) HEL and (**H**) KG-1 cells +/-5µM CHIR99021 and +/-20µg/mL RNaseA. All immunoblots shown are representative of 3 independent biological replicates.

To examine the subcellular location of the β-catenin:WT1 interaction we performed co-localisation studies using confocal laser scanning microscopy (CLSM). Using KG-1, K562 and HL60 cell lines, we observed WT1 is mainly a nuclear protein with some colocalisation with β-catenin (mainly cytosolic) during basal Wnt signalling. However, a significant increase in the colocalisation of β-catenin and WT1 signal is observed in the nucleus during stimulated Wnt signalling in all 3 cell lines (Figure 2A and B) including NB4 and HEL cells (Supplementary Figure S1D). To evaluate the clinical relevance of this protein interaction we examined β-catenin and WT1 expression across a panel of primary AML patient samples by immunoblotting (clinical details in Supplemental Table S1). We observed that approximately one third (9/30) of this cohort co-overexpressed both proteins relative to normal CD34^+^ cord blood (CB)-derived HSPC (Figure 2C) – a similar frequency of coexpression to that observed in myeloid cell lines (Figure 1A). We detected β-catenin expression in HSPC in keeping with its self-renewal role in this context,^11^ but WT1 was undetectable as expected given only 1.2% of the CD34^+^ HSPC pool are estimated to express this protein.^12^ Overall, we observed WT1 overexpression in around 47% (14/30) of our AML patient blast screen, consistent with previous estimates of high WT1 overexpression in this disease.^6,13^ As with the cell line correlation, there was no obvious association of this co-expression with any particular clinical characteristic. From this screen we performed a nuclear/cytosol fractionation on selected AML samples co-overexpressing both proteins and observed high levels of both proteins in the nucleus and cytoplasm, including the presence of the active (non-phosphorylated) form of β-catenin (Figure 2D). This would indicate the β-catenin:WT1 interaction has the potential to be cytoplasmic or nuclear in primary AML patient blasts as previously shown in HEL cells.^4^ We further used AML patient sample (#4) which expressed high levels of both proteins (and where ample viable cellular material was available) to perform a WT1 Co-IP and confirmed the β-catenin:WT1 interaction by immunoblot (Figure 2E). Finally, we performed a single tandem mass tag (TMT) labelled mass spectrometric (MS) analysis of the β-catenin interactome (using β-catenin Co-IP versus IgG Co-IP) in the highest β-catenin expressing AML patient sample (#1) and revealed a ∼70% enrichment of WT1 in the nucleus (Supplemental Table S2 and Supplementary MS data). Taken together, these data demonstrate for the first time that β-catenin and WT1 interact in both myeloid cell lines and primary AML samples.

**Figure 2.**
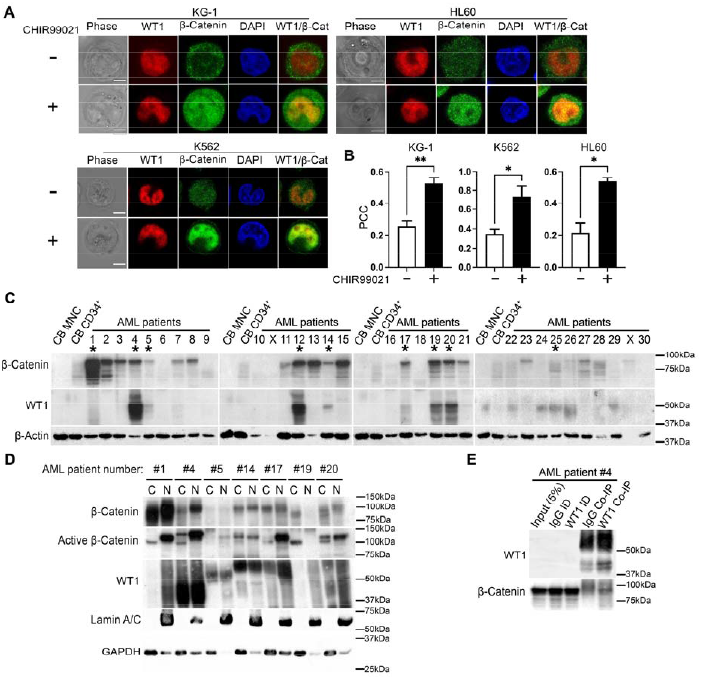
Colocalization and clinical relevance of β-catenin:WT1 interaction. (**A**) Representative CLSM Z-sections showing β-catenin and WT1 subcellular localisation in KG-1, K562 and HL60 cells +/-5µM CHIR99021. Phase (gray), WT1 (red), β-catenin (green), DAPI (blue) and merged WT1/β-catenin images are shown. Data shown are representative of 20 individual cells derived from 3 independent experiments, white scale bar indicates 5µm. (**B**) Correlation between β-catenin and WT1 signal in KG-1, K562 and HL60 cells +/-5µM CHIR99021, as determined by Pearsons Correlation Coefficient (PCC; -1= inverse correlation, 0= no correlation, +1= positive correlation). All data represents mean_l_±_l_1 s.d as deduced from a minimum of 3 independent experiments each containing a minimum of 20 cells per field. Statistical significance is denoted by \**P*<0.05 and \*\**P*<0.01 as deduced from a student’s t-test. (**C**) Immunoblot screen of 30 primary AML patient samples showing the relative level of β-catenin and WT1 protein; * denotes samples overexpressing both WT1 and β-catenin relative to levels in cord blood derived mononuclear cells (CB MNC) and CD34^+^ enriched fraction (CB CD34^+^) pooled from five independent cord blood samples. X = void sample as deduced from β-actin which assessed protein loading. (**D**) Immunoblot showing total β-catenin, active β-catenin and WT1 localization in selected AML patient samples co-overexpressing both β-catenin and WT1 (from initial screen). Lamin A/C and GAPDH indicate the purity/loading of the nuclear (N) and cytosol (C) fractions. (**E**) Immunoblot showing the level of β-catenin protein present in WT1 Co-IP performed from primary AML patient sample #4 of sample screen.

Given that both Wnt and WT1 signalling have been heavily implicated in AML we wanted to assess the potential signalling interplay between these two proteins in AML cells given crosstalk has been identified in other contexts.^14^ Using KG-1 cells (an AML cell line expressing both proteins that could tolerate WT1 knockdown) we knocked down WT1 in KG-1 cells using two different shRNA sequences and observed a consistent reduction in the β-catenin nuclear localisation capacity (Figure 3A). This corresponded with a significant reduction in basal, pharmacologically activated (CHIR99021), and naturally activated (rWNT3A) Wnt signalling output using the β-catenin activated reporter (BAR) system (Figure 3B, C and D). This supports previous studies demonstrating cooperation between WT1 and Wnt signalling in other systems.^15,16^

**Figure 3.**
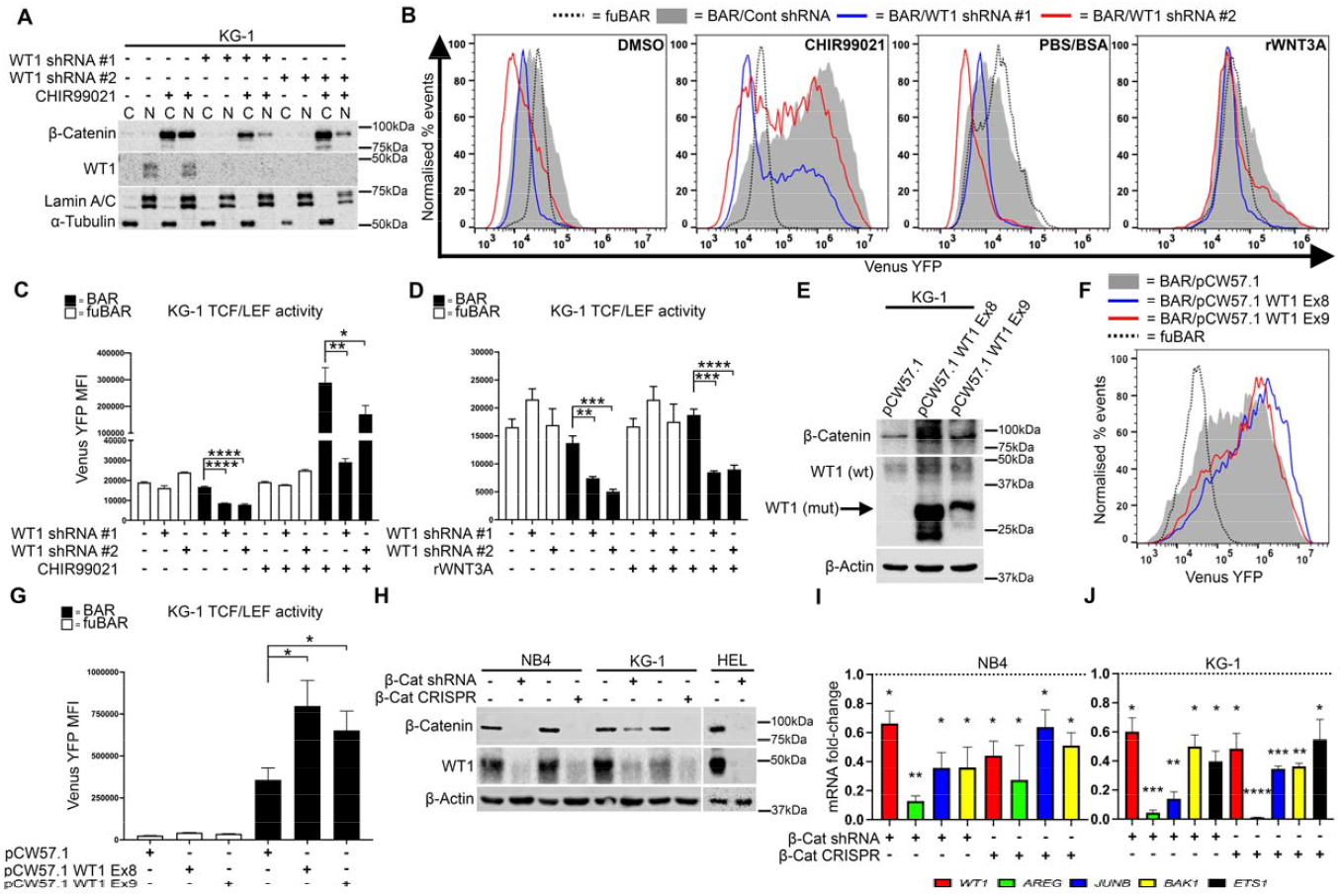
Crosstalk between Wnt and WT1 signalling in AML cells. (**A**) Immunoblot showing total β-catenin and WT1 subcellular localization in KG-1 cells lentivirally transduced with two different WT1 shRNAs +/-5µM CHIR99021. Lamin A/C and α-tubulin indicate the purity/loading of the nuclear (N) and cytosol (C) fractions respectively. (**B**) Representative flow cytometric histograms showing intensity of the TCF-dependent expression of Venus Yellow Fluorescent Protein (YFP) from the β-catenin activated reporter (BAR) reporter, or negative control ‘found unresponsive’ BAR (fuBAR; containing mutated promoter binding sites) in KG-1 cells +/-WT1 shRNA +/-5µM CHIR99021 or +/-1µg/mL rWNT3A. The fuBAR (dashed), non-targeting control shRNA (grey filled), and two WT1 shRNAs (blue or red) histograms are shown. Summary graphs showing the median fluorescence intensity (MFI) generated from the BAR/fuBAR in KG-1 cells +/-WT1 shRNA with (**C**) +/-5µM CHIR99021 or (**D**) +/-1µg/mL rWNT3A. (**E**) Immunoblot showing total β-catenin and WT1 protein in doxycycline treated KG-1 cells expressing empty pCW57.1 vector, or pCW57.1 containing inducible exon 8 and exon 9 truncating mutations of WT1. Arrow indicates position of mutant WT1 protein and β-actin was used to assess protein loading. (**F**) Representative flow cytometric histograms showing the MFI of fuBAR/BAR in doxycycline treated KG-1 cells expressing empty pCW57.1 vector, or pCW57.1 containing inducible exon 8 or exon 9 WT1 mutations following treatment with 5µM CHIR99021. The fuBAR (dashed), empty pCW57.1 (grey filled), pCW57.1 exon 8 mutant WT1 (blue) and pCW57.1 exon 9 mutant WT1 (red) histograms are shown. (**G**) Summary graph showing the MFI generated by BAR/fuBAR from doxycycline treated KG-1 cells expressing empty pCW57.1 vector, or pCW57.1 containing inducible exon 8 or exon 9 WT1 mutations following treatment with 5µM CHIR99021. (**H**) Immunoblot showing total β-catenin and WT1 protein in KG-1, NB4 and HEL cells following lentiviral transduction with either β-catenin shRNA or CRISPR/Cas9 targeting β-catenin alongside matched respective controls. β-Actin was used to assess protein loading. Summary graph showing the fold change in WT1 target gene mRNA expression as assessed by qRT-PCR in (**I**) NB4 and (**J**) KG-1 cells expressing either β-catenin shRNA or CRISPR/Cas9. Fold change is relative to matched respective controls (dashed line) and overall expression was normalized to the housekeeping gene β-actin (*ACTB*). All data represents mean_l_±_l_1 s.d (*n*_l_=_l_3). Statistical significance is denoted by \**P*<0.05, \*\**P*<0.01, \*\*\**P*<0.001 and \*\*\*\**P*<0.0001 as deduced from a student’s t-test. Data shown are representative of 3 independent biological replicates.

WT1 mutations are frequent in AML, presenting in ∼10% of cases but the impact on Wnt/β-signalling has not previously been investigated.^6^ Using doxycycline (DOX)-inducible mutant WT1 expression constructs (kind gift of Constanze Bonifer)^5^ we examined the impact of WT1 mutations on β-catenin expression and TCF activity using frequently reported WT1 mutations. These variants include frameshift mutations to exons 8 and 9, which truncate the protein at different Zn^2+^-finger domains.^6^ The presence of these mutations has recently been found to significantly increase the growth and clonogenicity of AML cells as well as decreasing apoptosis.^5^ Using KG-1 cells we confirmed the expression of the truncated WT1 mutant proteins following DOX exposure, with the exon 8 mutant expressed more abundantly than the exon 9 mutant (Figure 3E). The presence of both mutations resulted in increased expression of the endogenous wild type WT1 as reported previously,^17^ and also a concomitant increase in total β-catenin expression (Figure 3E). Examination of Wnt signaling output using the BAR reporter showed that both WT1 mutants significantly augmented TCF activity (Figure 3F and G). These are the first studies to examine Wnt signalling in the context of WT1 mutations in AML and suggest the presence of WT1 mutations in AML cells can augment Wnt signalling activation.

To complement these studies, we also performed the reciprocal experiments to examine the effect of β-catenin knockdown on WT1 expression and signalling activity. Using both shRNA and CRISPR/Cas9 we successfully reduced β-catenin in AML cell lines able to tolerate its loss; both KG-1 and NB4 (Figure 3H). Only a short-term shRNA approach was possible in HEL cells, due to the lethality of β-catenin loss in these cells. In all cell lines we observed a dramatic decrease in total WT1 protein level (Figure 3H) in response to β-catenin loss. Like β-catenin, WT1 protein is ubiquitinated and regulated by the proteasome,^18^ therefore we hypothesised that the resultant WT1 loss upon β-catenin reduction might be a result of β-catenin protecting WT1 from proteasome-mediated degradation. However, treatment of both NB4 and KG-1 cells harbouring a β-catenin knockdown with the proteasome inhibitor MG132 failed to restore WT1 protein level, and instead reduced stability further (Supplemental Figure S1E). The loss of WT1 following proteasome inhibition has been reported previously following the treatment of myeloid cells with bortezomib which targeted WT1 transcript.^19,20^ Finally, to assess the impact of β-catenin on WT1 signalling activity we examined the mRNA expression of a panel of previously identified WT1 target genes using qRT-PCR including *WT1, AREG, JUNB, BAK1 and ETS-1*^7^ which were previously validated in KG-1 cells (Supplemental Figure S1F). In both NB4 and KG-1 cells we observed a significant reduction in all WT1 target genes assessed (except for *ETS1* in NB4 cells; data not shown), including *WT1* mRNA itself, upon either shRNA or CRISPR/Cas9 mediated β-catenin knockdown (Figure 3I and J). This suggests β-catenin mediated regulation of WT1 expression is at least in part transcriptionally driven. WT1 or its targets have not previously been identified as direct Wnt target genes (https://web.stanford.edu/group/nusselab/cgi-bin/wnt/target_genes), however very few such studies have been performed in a hematopoietic context, and our original study did not detect the β-catenin:WT1 interaction in colorectal cancer cells meaning this association could be highly context dependent.^4^ To our knowledge, this is the first report of β-catenin mediated regulation of WT1 expression/activity in AML. Like

β-catenin,^1,2^ WT1 is overexpressed in AML where it confers inferior prognosis,^5,6^ and also similar to β-catenin,^3,21^ WT1 cooperates with common genetic aberrations in AML such as t(8;21) *RUNX1::RUNX1T1*^22^ and t(9;11) *ML::AF9*^18^ to promote leukemogenesis. The results of this study, and previous studies suggesting functional overlap, raise the intriguing possibility of cooperation between these two frequently dysregulated proteins in AML. Such findings could be important for informing novel therapeutic strategies for targeting these oncoproteins in myeloid malignancies.

## Supporting information

Supplementary MS data for AML patient #1

## Acknowledgements

This work was funded by a Kay Kendall Leukaemia Fund Junior Fellowship (RM; KKL1051) and a University of Sussex, School of Life Sciences PhD studentship (MW). We are grateful to Luke Ames, Dr Sandeep Potluri and Professor Constanze Bonifer for the kind sharing of inducible mutant WT1 expression vectors. Special thanks to Dr Paraskevi Diamanti (University of Bristol) for AML patient sample collection and Lizzy Hoole (Institute for Child Life &Health, UHBristol &Weston NHS Foundation Trust) for supplying clinical data. Thanks to the midwives and clinical research nurses at Brighton &Sussex University Hospitals NHS trust including Raquel Akieme and Caroline Humphreys, for the collection of human umbilical cord blood. We are indebted to the patients and their families who gave consent for their cells to be used for our research.

In loving memory of Dr Stephen Hare. A treasured colleague, researcher, teacher, friend, husband, and father taken too soon.

