## Supplementary data and methods for "Crosstalk between β-catenin and WT1 signalling activity in acute myeloid leukemia"

**Supplementary Table S1. Clinical characteristics of AML/MDS patient diagnostic/relapse samples used in this study**.

| **Patient no.** | **Age (at diagnosis)** | **Sex** | **WBC count (x10^9^/L)** | **Sample type** | **Secondary disease (Y/N)** | **Genetic information** | **Other clinical information** |
| --- | --- | --- | --- | --- | --- | --- | --- |
| 1 | 76 | F | 389 | LP | N | Normal karyotype, NPM1^+^, FLT3^+^ | n/a |
| 2 | 4 | M | n/a | BM | n/a | n/a | n/a |
| 3 | 10 | F | n/a | BM | n/a | n/a | Deceased |
| 4 | n/a | n/a | >200 | PB | Y | n/a | Post-allogeneic transplant. M0/1 (previously diagnosed with M3 10 years previously) |
| 5 | n/a | n/a | n/a | n/a | n/a | n/a | n/a |
| 6 | 17 | M | n/a | BM | Y | n/a | Post-BMT for AML following 2 relapses. Deceased |
| 7 | 6 | M | 70.7 | n/a | Y | n/a | Secondary to Ewings Sarcoma. Myelomonocytic morphology. Deceased. |
| 8 | 14 | F | 7.6 | BM | N | MLL rearrangement. Karyotype: 46,XX,ins(10;11)(q11.2;q23.1q23.3).ish ins(10;11)?inv(11)(q23.3)(5’MLL+)(q23.1)(3’MLL+) | BMT for high-risk AML. CD33+, MPO+, CD34-, CD117+, TdT+, CD64+, CD11c+, CD15+, CD11b+, NG2+ |
| 9 | 24 | M | 256 | BM | N | 46,XY,inv(16)(p13q22)[16].ish inv(16)(p13)(MYH11+,CBFB+)(q22)(CBFB+,MYH11+)[5] | M4, relapsed, deceased. |
| 10 | 17 | M | 294 | BM | N | 46,XY[20] | Alive, complete remission. |
| 11 | 64 | F | 13.3 | BM | N | Normal karyotype | AML with underlying MDS like changes |
| 12 | 13 | F | 91.9 | BM | Y |  | High Risk. Relapsed AML secondary to Rhabdoid tumour. Mixed cellular infiltrate consisting of predominantly monocyte/macrophages (CD14+, CD11c+, CD64+, CD34-, CD117-). Deceased |
| 13 | 4 | F | 2.6 | BM | N | Normal karyotype, NPM1^+^ (exon 12), FLT3^-^ | MRD neg. CD13+, CD33+, CD34+, CD117+, MPO+ |
| 14 | 15 | F | 6.5 | BM | N | MLL (KMT2A) rearrangement, t(10;11)(p11-p14,q23), MLL-MLLT10 | MRD detected post treatment course 1 |
| 15 | 4 | M | 3.2 | BM | N | t(10;11)(p11.2;q23) KMT2A-MLLT10. Cytogenetically cryptic. KMT2A ex8-MLLT10 ex9 or KMT2A ex9-MLLT10 ex10 fusion detected. NPM1^-^FLT3^-^ | CD13-, CD33+, CD34-, CD117+/-, CD11c+, CD64+, CD14-, NG2+. High risk cytogenetics. BMT. MRD neg post course 1+2. |
| 16 | 14 | F | 14.7 | PB | N | t(8;21)(q22;q22) RUNX-RUNX1T1. 46,XX,der(8)?del(8)(q11.2q21.3)?dup(8)(q24.3q21.3)?ins(8;21) (q22;q22.12q22.3),der(21)?ins(8;21)(q22;q22.12q22.3)[10] | CD13+, CD33+, CD34+, CD117+, DR+, MPO+, CD19+. MRD negative post treatment course 1, 2 and EOT |
| 17 | 5 | M | 1.6 | BM | N | n/a | M5. Deceased. |
| 18 | 14 | F | 2.1 | BM | N | 45,XX,-7,add(11)(p11.2)[9]/46,XX[1] | 22% Myeloid blasts (CD117+, CD34+/-, CD33+, CD13+, MPO-) + 10% immunophenotypically mature monocytes (CD11c+, CD64+, CD117+) BMT |
| 19 | 8 | F | n/a | BM | N | n/a | M4/5. Deceased. |
| 20 | 7 | F | 34.4 | BM | Y | MLL rearrangement t(9;11) | M5a morphology. BMT following relapse. Deceased |
| 21 | 16 | F | 20.6 | BM | N | Normal karyotype 46,XX[20] | CD33+, MPO+, CD34+, CD117+, CD13+, CD14-, CD7+, CD45 weak, CD11c+, TdT- |
| 22 | 7 | M | 3.5 | BM | N | t(8;21)(q22;q22) RUNX-RUNX1T1  45,X,-Y,t(8;21)(q22;q22)[9]/46,XY[1] | 8% myeloid blasts present (CD13+, CD33+, CD34+, CD117+, MPO+) |
| 23 | 11 | M | n/a | BM | N | n/a | BMT for MDS. Deceased. |
| 24 | 13 | F | 14.7 | BM | N | Karyotype: 46,XX,der(8)?del(8)(q11.2q21.3)?dup(8)(q24.3q21.3)?ins(8;21)(q22;q22.12q22.3),  der(21)?ins(8;21)(q22;q22.12q22.3)[10]  CD13+, CD33+, CD34+, CD117+, DR+, MPO+, CD19+.; AML | Only 2 megakaryocytes seen. Erythropoiesis reduced. Prominent eosinophils and eosinophil precursors - c. 12%. No significant monocytoid population. Densely infiltrated with myeloid blasts.  Analysis showed an abnormal female clone with a derivative chromosome 8 from a variant 8;21 rearrangement. This abnormality is consistent with a diagnosis of AML (WHO 2008 subtype: AML with t(8;21)(q22;q22); RUNX1-RUNX1T1) and is reported in association with a favourable prognosis.  Alive, complete remission. |
| 25 | 2 | M | 2.4 | BM | N | t(9;11)(p22;q23), t(11;21)(q23;q8) | n/a |
| 26 | 4mo | M | 5.1 | BM | N | t(9;11) | BMT |
| 27 | 7 | F | n/a | BM | N | MPAL, 46XX, del5q, abnormal 21 | n/a |
| 28 | 7mo | F | 168.6 | BM | N | Karyotype: 47,XX,+21[5]/46,XX[5] nuc ish(CBFA2T3,GLIS2)X3(CBFA2T3 con GLIS2x2)[92/150]  + RUNX1. | BMT 24/12/20 due to high-risk genetics.  CD33+, CD34+, CD117+, MPO+, DR-, CD13+  Alive, in remission.  Received Gemtuzumab Ozogamycin as part of Myechild trial treatment. |
| 29 | 5 | F | 29.5 | BM | N | Karyotype: 45,X,-X,t(8;21)(q22;q22.1)[8]/46,XX[2] | 7% myeloid blasts (CD13+, CD33+, CD34+, CD117+, MPO+)  Alive, in remission.  Received 2 doses of Gemtuzumab Ozogamycin as part of Myechild trial treatment |
| 30 | 76 | F |  | BM |  |  | D45X Polycythaemia vera. M99503, transformation in to MDS. |

BM = Bone marrow

PB = Peripheral blood

LP = Leukapheresis

MRD = Minimal residual disease

BMT = Bone marrow transplant

AML= Acute myeloid leukemia

MDS= Myelodysplastic syndrome

MPAL= Mixed phenotype acute leukemia

MLL = *Mixed-lineage leukemia*

NPM1 = *Nucleophosmin*

FLT3 = *Fms-like tyrosine kinase 3*

RUNX1 = *Runt-related transcription factor 1*

GATA2 = *GATA Binding Protein 2*

WHO = World Health Organisation

n/a = not available

**Supplementary Table S2. Enrichment values for β-catenin and WT1 following a TMT-labelled assessment of AML Patient #1 β-catenin interactome by mass spectrometry.**

| **Protein** | **Score** | **Coverage** | **# Proteins** | **# Unique peptides** | **# Peptides** | **PSM** | **Cyt fold change (vs IgG)** | **Nuc fold change (vs IgG)** |
| --- | --- | --- | --- | --- | --- | --- | --- | --- |
| **β-Catenin** | 535.53 | 64.08 | 10 | 33 | 40 | 175 | 8.03 | 4.159 |
| **WT1** | 2.51 | 4.49 | 5 | 1 | 1 | 1 | 0.889 | 1.74 |

**Score:** Displays the protein score, which is the sum of the scores of the individual peptides.

For Sequest results, the score is the sum of all peptide Xcorr values above the specified score threshold. The score threshold is calculated as follows:

0.8 + peptide_charge × peptide_relevance_factor

where peptide_relevance_factor is an advanced parameter of the SEQUEST or Sequest HT node in the “Protein Scoring Option” category with a default value of 0.4. For each spectrum, only the highest-scoring match is used.

For each spectrum and sequence, the Proteome Discoverer application uses only the highest scored peptide. When it performs a search using dynamic modifications, one spectrum might have multiple matches because of permutations of the modification site.

For Mascot results, the score is:

– Standard score, which is the cumulative protein score based on summing the ion scores of the unique peptides identified for that protein. If a peptide was redundantly identified, only the highest-scoring peptide is used.

–or–

– MudPIT score, which is the sum of the “excess of ions” score over the homology or identity threshold for each spectrum plus the average threshold. For MudPIT scoring, the score for each peptide is not its absolute score but the amount that it is above the threshold. Therefore, peptides with a score below the threshold do not contribute to the score. For each peptide, the threshold is the homology threshold, if it exists; otherwise, it is the identity threshold. By default, the Proteome Discoverer application automatically switches between the standard and the MudPIT score to calculate the protein score in the Mascot node results. It automatically uses the MudPIT score when the number of queries divided by the number of FASTA database entries exceeds 0.001.

**Coverage:** Displays by default the percentage of the protein sequence covered by identified peptides

**Proteins:** Displays the number of identified proteins in the protein group of a master protein

**Unique Peptides:** Displays the number of peptide sequences unique to a protein group

**Peptides:** Displays the number of distinct peptide sequences in the protein group

**PSMs:** Displays the total number of identified peptide sequences (peptide spectrum matches) for the protein, including those redundantly identified

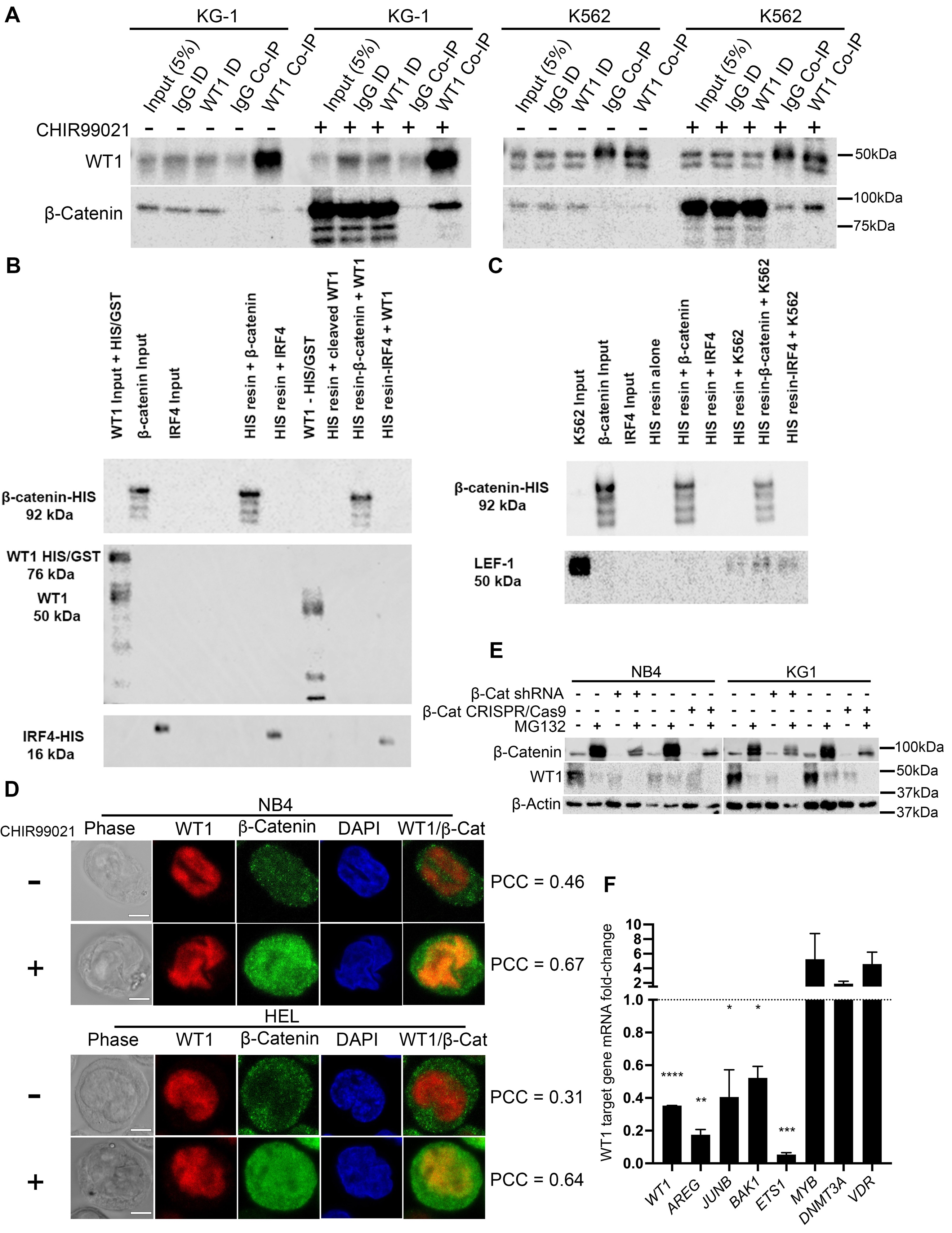

**Supplementary Figure S1.** (**A**) Immunoblots showing the level of β-catenin protein present in WT1 Co-IPs derived from KG-1 and K562 under basal (DMSO)or activated (5µM CHIR99021) Wnt signalling conditions. ID= immunodepleted lysate. (**B**) Immunoblot showing purified WT1 (HIS/GST tagged, and non-tagged) abundance in β-catenin-HIS or IRF4-HIS (negative control) resin columns. (**C)** Immunoblot showing level of β-catenin or positive control LEF-1 (derived from whole cell K562 lysates) present in HIS resin β-catenin and IRF4 containing lanes. LEF-1 is a known interactor of β-catenin and confirms recombinant β-catenin protein can still bind established partners. (**D**) CLSM Z-sections showing β-catenin and WT1 subcellular localisation in NB4 and HEL cells +/- 5µM CHIR99021. Phase (gray), WT1 (red), β-catenin (green), DAPI (blue) and merged WT1/β-catenin images are shown alongside Pearsons Correlation Coefficients (PCC; -1= inverse correlation, 0= no correlation, +1= positive correlation) indicating degree of overlap between β-catenin and WT1 signal. White scale bar indicates 5µm. (**E**) Immunoblot showing protein level of β-catenin and WT1 in NB4 and KG-1 cells +/- β-catenin shRNA, +/- β-catenin CRISPR/Cas9 following 16 hours incubation with 1µM proteasome inhibitor MG132. β-Actin detection indicates protein loading. (**F**) Summary graph showing fold change in mRNA expression of genes previously identified as WT1 target genes, in KG-1 cells by qRT-PCR. Fold change is in response to knockdown of WT1 using WT1 shRNA relative to expression in non-targeted shRNA control (dashed line). Expression was normalised to the housekeeping gene β-actin (*ACTB*). All data represents mean ± 1 s.d (*n* = 3). Statistical significance is denoted by **P*<0.05, ***P*<0.005, ****P*<0.0005 and *****P*<0.0001 as deduced by a one-sample t-test.

**Methods**

*Primary samples*

Bone marrow, peripheral blood or leukapheresis samples from patients diagnosed with AML/MDS (Clinical information in Supplementary Table S1) were collected in accordance with the Declaration of Helsinki and with approval of University Hospitals Bristol and Weston NHS Foundation Trust and London Brent Research Ethics Committee. Human cord blood was obtained following informed consent from healthy mothers at full-term undergoing elective caesarian sections at the Royal Sussex County Hospital, with approval from Brighton & Sussex University Hospitals NHS (BSUH) trust, the East of England – Essex Research Ethics Committee, Human Research Authority and Health and Care Research Wales (18/EE/0403). From all primary samples mononuclear cells (MNC) were separated using Ficoll-Hypaque (Merck Millipore, Dorset, UK) and samples with ≥80% viability included in the study. CD34^+^ cells were derived as previously described^1^ from cord blood MNC and enriched to >95% purity using MiniMACS (Miltenyi Biotec, Surrey, UK) according to the manufacturer's instructions.^2^

*Cell culture and β-catenin stabilization*

The myeloid cell lines K562, HL60, HEL, U937, PLB-985, NOMO1, OCI-AML3, MOLM-13, EOLI, ML-1, THP-1 (ECACC, Salisbury, UK), MV4-11, NB4 and MonoMac6 (DSMZ-German, Braunschweig, Germany) were cultured, and β-catenin stabilised using the GSK-3β inhibitor CHIR99021 (Merck Millipore) or recombinant murine WNT3A as previously described.^2^

*WT1 co-immunoprecipitation and immunoblotting*

250µL of WT1 antibody (Clone 89, gift of Prof Stefan Roberts, University of Bristol, UK) or 5μg of rabbit IgG (Merck Millipore, Dorset, UK) was crosslinked to Protein G Dynabeads (Thermofisher Scientific, Waltham, MA) and Co-IPs performed as previously described,^2^ but for the inclusion of Co-IP buffer containing 20μg/mL RNase A (Thermofisher Scientific). Samples were prepared and immunoblotted as previously^2^ with antibodies to β-catenin (Clone 14, Becton Dickinson (BD), Oxford, UK), active β-catenin (Clone 8E7, Merck Millipore) WT1 (Clone CAN-R9(IHC)-56-2, Abcam, Oxford, UK), GAPDH (Clone 1E6D9, Proteintech, Manchester, UK) and β-actin (Clone AC-15, Merck Millipore). Densitometry for WT1 and β-catenin expression was performed using ImageJ software version 1.5.2 (Tree Star Inc., Ashland, OR) normalising to the β-actin density present within each respective sample.

*Agarose gel electrophoresis*

Reactions were heated at 37°C for 5 minutes and electrophoresed on a 1% (*w*/*v*) agarose gel (Thermofisher Scientific) by dissolving the required amount of agarose powder in 1x Tris-Borate-EDTA (TBE; Thermofisher Scientific) and heating until a homogenous solution was achieved. After the solution had cooled 2.5μL of gel red (Biotium, San Francisco, CA) was added, poured, and left to set at room temperature. Samples were prepared with 6X gel loading buffer (New England Biolabs, Hitchin, UK) and electrophoresed at 75V for 30 minutes.

*Immunofluorescence*

DMSO/CHIR99021 treated cell lines were fixed in 2% paraformaldehyde (Merck Millipore) for 20 minutes at room temperature. Cells were quenched (1 x PBS, 100mM glycine), followed by permeabilization (1 x PBS, 0.1% Triton TX100) and washing. Fixed and permeabilised cells were resuspended in staining buffer (1xPBS, 0.5% BSA) containing 10μg/mL antibodies to WT1 (Clone CAN-R9(IHC)-56-2) and β-catenin (Clone 14) for 30 minutes at room temperature. Following washing, cells were resuspended in 1mL of staining buffer containing both 4µg/mL Alexa488 conjugated goat anti-mouse and Alexa647 conjugated goat anti-rabbit secondary antibodies (Invitrogen, Paisley, UK) for 30 minutes in the dark at room temperature. Finally, cells were resuspended in staining buffer containing 10µg/mL DAPI (Thermofisher Scientific) for 5 minutes at room temperature. Images were captured using the resonant scanning head of a Zeiss LSM880 confocal microscope with a 40X oil immersion objective and assisted by Zen software version 3.4. Post-acquisition editing of images, generation of overlays and execution of colocalization analyses were done using ImageJ software and plugins.

*Nuclear/Cytoplasmic fractionation*

Performed as previously described.^2^

*β-Catenin interactome mass spectrometry analyses*

β-catenin antibody crosslinking, β-catenin Co-IP, Tandom Mass Tag (TMT)-labelling and mass spectrometry analyses were executed and analysed as previously described.^2^

*Lentiviral transduction*

Cell lines were lentivirally-transduced with the β-catenin-activated reporter (BAR) or mutant ‘found unresponsive’ control (fuBAR) system as previously described.^3^ Cells were lentivirally transduced with either human WT1 shRNAs (TRC0000-040063, -009873; MISSION® Merck Millipore), β-catenin shRNA (Plasmid #18803; Addgene, Cambridge, MA) or β-catenin CRISPR/Cas9 (VB181017-1120bsy; VectorBuilder, Neu-Isenburg, Germany). For WT1 mutation studies, KG1 cells were transduced with either empty pCW57.1 vector (Addgene), or pCW57.1 vector encoding inducible frameshift mutations in either exon 8 or exon 9 which truncated WT1 at different zinc finger domains (kind gift of Constanze Bonifer, University of Birmingham, UK) and were induced using 2µg/mL doxycycline (Merck Millipore).^4^ Cells were transduced with the relevant non-targeting scrambled shRNA or CRISPR/Cas9 vectors as controls.

*Assessment of TCF reporter and flow cytometry*

TCF activity was assessed using the BAR system as previously described.^3^ Multi-parameter flow cytometric measurements were acquired using an Accuri C6 in conjunction with C sampler software v1.0.264.21 (BD). Post-acquisition analyses were performed using FlowJo v10.8.0 (BD). Threshold for TCF reporter fluorescence was set using matched-controls expressing mutant ‘found unresponsive’ fuBAR. Cell viability was assessed using 7-Aminoactinomycin D (7AAD; Thermofisher Scientific).

*qRT-PCR*

Extraction of total RNA was performed by lysing 5x10^6^ cells in 1mL of Trizol (Thermofisher Scientific), 200µL of chloroform (Thermofisher Scientific) was added and samples spun at 17,000 x *g* for 15 minutes at 4°C to ensure separation with RNA in the upper aqueous phase. This was removed and mixed with 500µL isopropanol (Thermofisher Scientific), centrifugation was repeated, and the pellet was washed in 80% (*v*/*v*) ethanol (Merck Millipore). cDNA synthesis was performed using 2µg of total RNA using a high-capacity cDNA reverse transcription kit (Thermofisher Scientific). Forward and reverse primers (sequences below) for target genes *WT1*, *AREG*, *JUNB*, *BAK1* and *ETS1* (Merck Millipore) were prepared with cDNA and Go-Taq SYBR (Promega, Wisconsin, USA) in a MicroAmp 96 fast optical plate (Thermofisher Scientific) with an optical film (Thermofisher Scientific) placed on top. The plate was analysed using the StepOnePlus QPCR machine (Thermofisher Scientific), and target gene expression levels were normalised to non-targeted control cells and β-actin using StepOne software v2.3 (Thermofisher Scientific).

**Supplementary Table 3. Sequences for forward and reverse primers used in this study.**

| **Gene Name** | **Direction** | **Tm°** | **Length (bp)** | **5’ -> 3’ Sequence** |
| --- | --- | --- | --- | --- |
| *ACTB* | Forward | 66.1 | 22 | TTGTTACAGGAAGTCCCTTGCC |
| *ACTB* | Reverse | 67.2 | 22 | ATGCTATCACCTCCCCTGTGTG |
| *WT1* | Forward | 60 | 20 | CACAGCACAGGGTACGAGAG |
| *WT1* | Reverse | 60 | 20 | CAAGAGTCGGGGCTACTCCA |
| *AREG* | Forward | 50 | 20 | TGGATTGGACCTCAATGACA |
| *AREG* | Reverse | 50 | 20 | ACTGTGGTCCCCAGAAAATG |
| *ETS1* | Forward | 60 | 23 | AAACTTGCTACCATCCCGTACGT |
| *ETS1* | Reverse | 60 | 22 | ATGGTGAGAGTCGGCTTGAGAT |
| *JUNB* | Forward | 60 | 21 | TGGTGGCCTCTCTCTACACGA |
| *JUNB* | Reverse | 60 | 17 | GGGTCGGCCAGGTTGAC |
| *VDR* | Forward | 60 | 20 | CTGACCCTGGAGACTTTGAC |
| *VDR* | Reverse | 60 | 19 | TTCCTCTGCACTTCCTCAT |
| *BAK1* | Forward | 60 | 21 | GTTTTCCGCAGCTACGTTTTT |
| *BAK1* | Reverse | 60 | 22 | GCAGAGGTAAGGTGACCATCTC |
| *DNMT3A* | Forward | 60 | 20 | CCGATGCTGGGGACAAGAAT |
| *DNMT3A* | Reverse | 60 | 20 | CCCGTCATCCACCAAGACAC |
| *MYB* | Forward | 60 | 21 | GAAAGCGTCACTTGGGGAAAA |
| *MYB* | Reverse | 60 | 23 | TGTTCGATTCGGGAGATAATTGG |

*Recombinant protein expression*

Full length WT1 and β-catenin were cloned into pET479b and pET47b vectors respectively. WT1 was overexpressed with an N-terminal GST HIS_6_ tag in *Escherichia Coli* BL21 cells (Merck Millipore) at 18°C overnight and β-catenin was overexpressed with an N-terminal HIS_6_ tag in BL21 cells at 37°C for 4 hours. Cells were induced with 0.5mM IPTG (Merck Millipore). Cells were resuspended in 10mL of lysis buffer (20mM HEPES pH 7.5 (Thermofisher Scientific), 0.5M NaCl, 0.1mM ZnCl_2_ (Arcos Organics part of Thermofisher Scientific, Oxford, UK), 12% glycerol and 2mM TCEP (Thermofisher Scientific) or 10mL of β-catenin lysis buffer (20mM HEPES pH 7.5, 250mM NaCl, 10% glycerol and 2mM TCEP). Both lysis buffers contained an EDTA-free containing complete™ Mini Protease-Inhibitor Cocktail (Sigma Aldrich). Lysates were incubated on ice for 30 minutes and 2500U/mL of DNase (Thermofisher Scientific) was added. Lysates were sonicated (35% amplitude, 2.5 minutes (5 seconds on, 10 seconds off) per 10mL culture) and then centrifuged at 20,000 x *g* at 4°C for 30 minutes. The supernatant was removed and kept on ice. Proteins were purified using an affinity Econo-column (Bio-Rad) with specific resin to the GST (Generon, Slough, UK) or HIS tag (Expedeon, Oxford, UK) expressed by the plasmid in specific purification buffer with 5-500mM Imidazole (Arcos Organics, Oxford, UK) gradient. Optimal affinity column fractions were pooled and concentrated at 4000 x *g* for 20 minute intervals using Vivaspin concentrators (Sartorius, Surrey, UK) with an appropriate molecular weight cut off (20-100kDa depending on protein) to a final volume of 500μl and loaded onto a HiLoad 10/300 Superdex 200 gel filtration column (GE, Sigma Aldrich) pre-equilibrated with either WT1 SEC buffer (20mM HEPES pH 7.5, 200mM NaCl, 10% glycerol and 0.1mM ZnCl_2_ made to 1L with water) or β-catenin SEC buffer (20mM Tris pH 8, 150mM NaCl and 2mM TCEP made to 1L with water) connected to an AKTA purifier FPLC system (Cytiva, Marlborough, Massachusetts, USA). Fractions were eluted using the protein-specific SEC buffer at a flow of 0.45mL/min, monitoring protein elution by absorbance at 280nm and collecting 250μl fractions in to a 48 deep well plate. To cleave the HIS/GST tag from WT1, the eluted fractions were pooled and subjected to protease HRV3c 9 (gift from Mancini lab) digestion overnight at 4°C in WT1 purification buffer.

*HIS pull-down*

200μl of coHIS resin (50% slurry) was pre-equilibrated by washing three times with 500μl of pull-down buffer (20mM HEPES pH 7.5, 150mM NaCl and 2mM TCEP). 50μg/mL of β-catenin in pull-down buffer was incubated with tagged specific resin for 1 hour at 4°C under agitation to allow specific binding of the tagged over-expressed protein to the resin. The resin was pelleted by centrifugation at 800 x *g*, 4°C for 5 minutes and any unbound protein was removed, the resin was then washed three times with 1mL of pull-down buffer. 50µg/mL of reciprocal WT1 protein or 1mg of total cell lysate was incubated for 1 hour at 4°C under agitation, the resin was pelleted and washed three times.

*Statistics*

Statistical analyses were performed using GraphPad Prism v9.2.0 (GraphPad Software Inc., San Diego, CA). Correlation was assessed using a Spearman’s Rank correlation coefficient (R). Significance of difference was assessed using a one-sample or Students’ *t* test with the threshold for significance set at p<0.05 and data represents mean ±1 SD derived from three independent biological replicates.
